# Coordinate post-transcriptional regulation by microRNAs and RNA binding proteins is critical for early embryonic cell fate decisions

**DOI:** 10.1101/2024.10.02.616359

**Authors:** Carolyn Sangokoya, Robert Blelloch

## Abstract

Post-transcriptional control by RNA binding proteins (RBPs) and microRNAs play central roles in mRNA stability and translation (1). However, little is known about how RBPs and microRNAs coordinate in developmental time to regulate cell fate. Here, we show that coordinate RBP and microRNA control of a single transcript, Profilin 2 (Pfn2), is essential for differentiation of embryonic stem cells (ESCs) into the primary germ layer lineages. The Pfn2 3’untranslated region has a binding site for Iron Regulatory Proteins and a nearby binding site for ESC enriched microRNAs (2,3). Deletion of this microRNA site leads to increased PFN2 and reduced FGF signaling during pluripotency transition prior to germ layer formation (4). In contrast, deletion of the iron response element leads to decreased PFN2 and a Wnt signaling defect, reduced nuclear beta-catenin, and a subsequent block in mesendodermal lineages during early germ layer formation. The choreographed microRNA-IRE axis of control on the Pfn2 transcript is essential for two key signal transduction steps during ESC differentiation.

## Introduction

Mouse embryonic stem cells (ESCs) have the remarkable ability to self-renew indefinitely while retaining the potential to make all downstream lineages of adult body including the germline. To do so, they are typically grown in the presence of LIF and two inhibitors (2i) toward MEK (PD0325901) and GSK3 (CHIR99021) (5). MEK inhibition blocks FGF signaling while GSK3 inhibition promotes canonical Wnt signaling. GSK3 represses canonical signaling by phosphorylating the secondary messenger beta-catenin which directs it to the proteosome for degradation (6–8). ESCs grown under these conditions are called naïve ESCs and are transcriptionally similar to the *in vivo* pluripotent epiblast cells of the peri-implantation embryonic day 4.5 (E4.5) blastocyst (9). Upon removal of LIF and 2i, the naïve ESCs initiate differentiation, first transitioning to a more epithelial-like state resembling the pluripotent epithelial cells of the post-implantation E5.5 egg cylinder stage embryo, also called the formative pluripotent state (10). The cells can be retained in formative-like state by inhibiting Wnt signaling, providing low level Activin signaling, and allowing for autonomous FGF signaling (11). Following release of Wnt signaling, these cells than initiate differentiation into somatic germ layers, recapitulating gastrulation in vivo. This differentiation can occur spontaneously *in vitro* in the form of embryoid bodies or in more directed fashion in presence of specific factors (12–14). These well-defined methods of recapitulating early embryo development in a dish allows for an unprecedented ability to understand fundamental molecular and cellular mechanisms driving cell fate.

Post-transcriptional regulation of mRNA stability and protein translation is thought to play a central role in cell fate decisions, although only a limited number of examples exist. One such example is the role of the embryonic stem cell-enriched family of microRNAs that promote pluripotency including the dedifferentiation of somatic cells to induced pluripotent stem cells (3,15,16). MicroRNAs are short non-coding RNAs that function by binding the 3’UTR of target mRNAs, which in turn leads to both destabilization of the RNA and the inhibition of their translation (17–21).

The miR-290 cluster of microRNAs, including miR-291-3p, miR-294, and mir-295, makes up greater than 70% of total microRNAs in ESCs (22). The miR-290 cluster shares it binding site with the miR-302 cluster, which includes miR-302a-3p, miR-302b-3p, and miR-302d-3p). This binding site (AGCACUU) binds microRNAs miR-291-3p/294-3p/295-3p/302-3p and will be referred to as the miR290/302 site. While expression of the miR-290 cluster dominates ESC pluripotency, peaking in the peri-implantation blastocyst at E4.5, expression of the miR-302 cluster begins with early differentiation in parallel with decreasing miR-290 (23). Expression of miR-290 and miR-302 clusters overlap from post-implantation to early-mid gastrulation (∼E5.5-E7.0), as the miR-290 cluster is downregulated, and miR-302 expression begins and persists until E9.5 (24,25). Double knockouts of the miR-302/miR-290 cluster microRNAs leads to arrest peri-gastrulation (23), demonstrating the critical roles of the evolutionarily conserved miR-290 and miR-302 clusters during early mammalian development.

One functional target of the miR-290 cluster in promoting pluripotency is the actin/dynamin binding protein Profilin-2 (PFN2) (3). Deeper analysis of PFN2’s role has shown that deletion of the only miR290/302 site in the 3’UTR of PFN2 transcript leads to upregulation of PFN2 mRNA and protein levels, which in turn inhibits endocytosis-directed FGF signaling and thus the transition from naïve to formative pluripotency (4). There is also an iron response element (IRE) in the 3’UTR of PFN2 (2). This element is less from than 100bp from the miR290/302 site and is conserved through humans (Fig. 1a). IREs are target sites for the RNA-binding proteins (RBPs) Iron Response Proteins (IRP) 1 and 2. In contrast to microRNAs, binding of IRPs to IRE sites in the 3’UTR of a transcript typically results in its stabilization (26).

**Figure 1.**
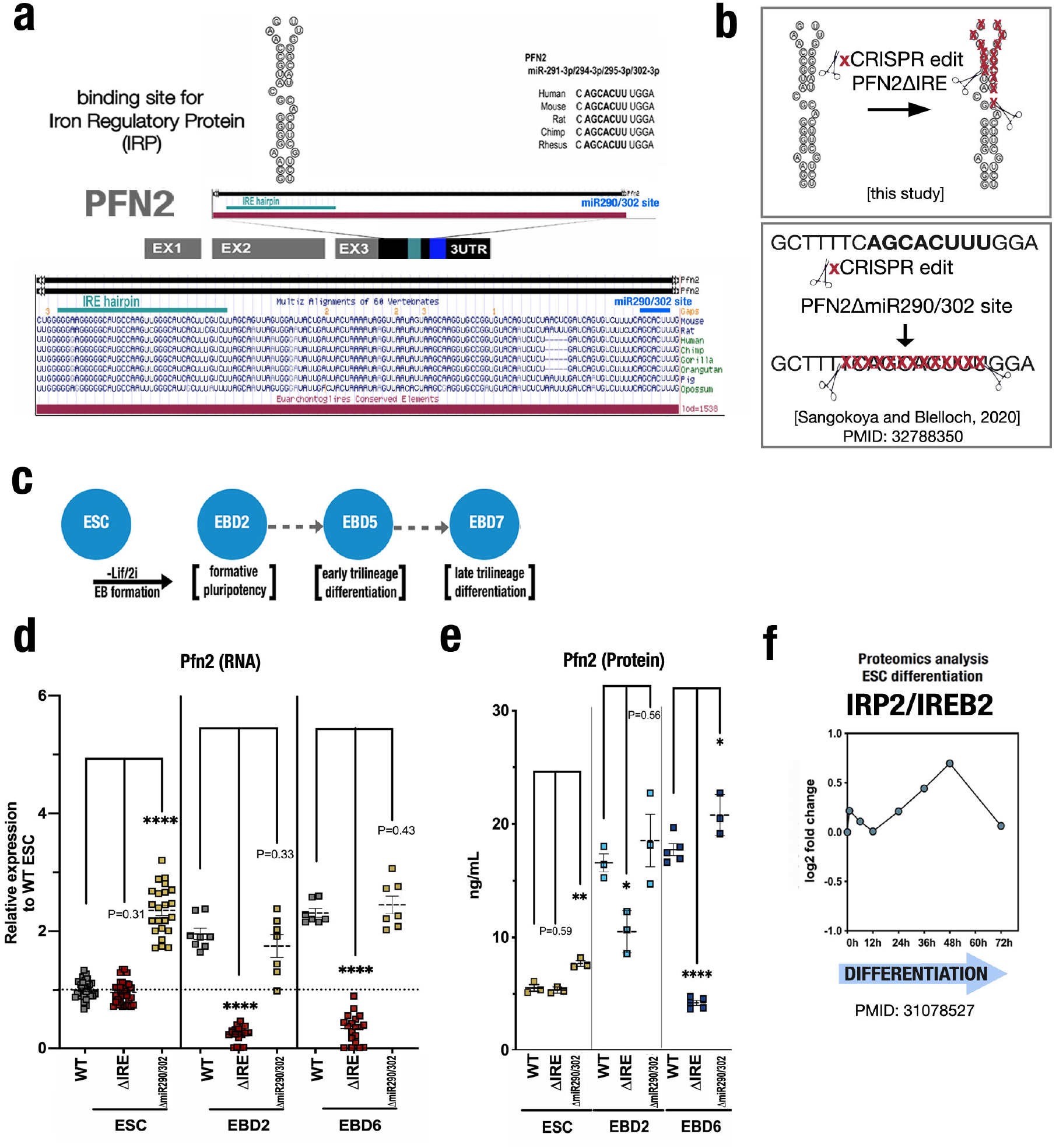
IRP/IRE site regulates PFN2 levels during ESC differentiation. **(a)** Schematic map of PFN2-3’UTR showing locations and evolutionary conservation of the neighboring IRP/IRE and Ago/miR-290 binding sites. **(b)** Representative schematic of CRISPR-edited PFN2-3UTR ΔIRE (top) and PFN2-3UTR ΔmiR290/302 (bottom) sequences with red x over deleted sequences **(c)** Schematic of (ESC) differentiation as embryoid bodies (EB) with descriptors of developmental stages at days 2, 5, and 7. **(d)** Expression analysis of Profilin-2 by qPCR in wild-type, PFN2-3UTR ΔIRE, and PFN2-3UTR ΔmiR290/302 mutants at naïve pluripotency (ESC): wild-type N=29; PFN2-3UTR ΔIRE N=38; PFN2-3UTR ΔmiR290/302 N=23; formative pluripotency (EBD2): wild-type N=8; PFN2-3UTR ΔIRE N=19; PFN2-3UTR ΔmiR290/302 N=7;, and early trilineage differentiation (EBD6): wild-type N=7; PFN2-3UTR ΔIRE N=19; PFN2-3UTR ΔmiR290/302 N=7 relative to wild-type ESC. Biological replicates for each cell line are shown across independent experiments. Error bars represent SEM. ****P<0.0001, unpaired two-tailed t test. **(e)** Absolute Profilin-2 protein levels (ng/mL) by ELISA in wild-type, PFN2-3UTR ΔIRE, and PFN2-3UTR ΔmiR290/302 mutants at naïve pluripotency (ESC): wild-type N=3; PFN2-3UTR ΔIRE N=3; PFN2-3UTR ΔmiR290/302 N=3;, formative pluripotency (EBD2): wild-type N=3; PFN2-3UTR ΔIRE N=3; PFN2-3UTR ΔmiR290/302 N=3;, and early trilineage differentiation (EBD6): wild-type N=5; PFN2-3UTR ΔIRE N=5; PFN2-3UTR ΔmiR290/302 N=3; relative to wild-type ESC. Biological replicates for each cell line are shown are shown across independent experiments. Error bars represent SEM. *P< 0.05, **P< 0.01, ****P<0.0001, unpaired two-tailed t test. **f)** Proteomics analysis result for IRP2 during ESC differentiation reported in cited study.

The presence of both a microRNA and an RBP site in close proximity on the 3’ UTR on the PFN2 target transcript raises the question of how they might be interacting to regulate ESC self-renewal and differentiation. Therefore, we deleted the IRE element in the PFN2 3’UTR and compared the consequences to loss of the miR290/302 site. Unlike loss of the miR290/302 site, disruption of the IRE site had no impact on PFN2 mRNA and protein levels in naïve ESCs. However, upon differentiation to the formative state and further into the three germ layers, disruption of the IRE element results in a reduction of PFN2 mRNA and protein levels. This reduction results in an inhibition of differentiation toward the three somatic germ layers, especially those of the primitive streak, mesoderm and endoderm. Additionally, this defect is associated with a decrease in Wnt signaling that is cell autonomous and is downstream of GSK3β directed degradation of beta-catenin. There is a decrease in beta-catenin in the nucleus, where it would normally be required to induce the gene expression program of mesendoderm differentiation. Therefore, these data show how the 3’UTR of a single gene can help direct the switch between pluripotency and the trilineage differentiation associated with gastrulation.

## Results

### Divergent post-transcriptional regulation of Profilin-2

To evaluate the role for the *Pfn2* 3’UTR iron response element (IRE) in ESC self-renewal and differentiation, we used CRISPR-Cas9 mutagenesis to delete this IRE site, generating PFN2-3UTR ΔIRE mutant ESCs. These cells were subcloned to produce a stable mutant cell line (Fig. 1b, Figure S1a-b). The PFN2-3UTR ΔIRE mutant ESCs were compared to matching wild-type control ESCs and previously produced PFN2-3UTR ΔmiR290/302 mutant ESCs (4). When cultured in naïve pluripotency media conditions (LIF+2i), the PFN2-3UTR ΔIRE mutant ESCs showed no significant difference in PFN2 mRNA and protein levels relative to corresponding wild-type cells (Fig. 1b-d). In contrast the PFN2-3UTR ΔmiR290/302 mutant ESCs showed elevated PFN2 mRNA and protein levels as previously described (4). To evaluate changes during ESC differentiation, we performed a time course of embryoid body (EB) differentiation (Fig 1b). Upon EB differentiation, wild-type ESCs show an approximately two-fold increase in Profilin-2 mRNA levels and three to four-fold increase in protein levels starting at day 2 (EBD2, representing the naïve to formative pluripotency transition) through day 6 (EBD6, representing germ layer specification) (Fig. 1c,d). This normal increase in Pfn2 protein with differentiation is significantly decreased in PFN2-3UTR ΔIRE mutant cells (Fig. 1c,d). In contrast, PFN2-3UTR ΔmiR290/302 cells show similar levels of PFN2 mRNA and protein as wild-type cells at EBD2 and EBD6. These findings uncover divergent roles for the miR290/302 and IRE sites in the 3’UTR in PFN2. The miR290/302 site suppresses PFN2 levels in naïve ESCs, while the IRE site is essential for the stability of the mRNA and upregulation of PFN2 protein upon ESC differentiation.

The iron regulatory proteins IRP1 and IRP2 are necessary for cellular iron homeostasis and can both bind IRE sites. IRP2 is the dominant of the two IRPs in terms of iron regulation (27) and is downregulated at the protein level when not serving as an RNA-binding regulator of iron homeostasis (26–30). Recent proteomic analysis during ESC differentiation highlights the peak level of IRP2 48 hours after differentiation, identical to the EBD2 timepoint in our studies (31) (Fig. 1f). Additionally, a recent IRP1/2 activity sensor that incorporates the sum of IRP1/2 activity was tested during ESC differentiation and also recapitulates this IRP activity peak at EB2/48 hours differentiation (32).

### Loss of the PFN2-3UTR IRE site inhibits ESC differentiation

To characterize the impact of the PFN2-3UTR IRE site on the formation of the three germ layers, we performed embryoid body differentiation of wild-type and PFN2-3UTR ΔIRE mutants at EBD6 followed by RNA-seq. Over 4000 genes were dysregulated in the mutants (1815 up, 2261 down, FDR <0.05) (Fig. 2a). Compared to wild-type cells, PFN2-3UTR ΔIRE mutants showed reduced expression of markers for all 3 germ layers including mesoderm (*Mixl1, Col4a1, Col4a2*), endoderm (*Pdgfra, Sox17, Hnf4a, Hnf1b*) and ectoderm (*Sox11, Pax6*), (Fig. 2a). In contrast, pluripotency markers were abnormally elevated. Gene ontology analysis showed enrichment for categories such as cell migration involved in gastrulation, endoderm formation, and mesoderm formation (Fig. 2b). A closer look at well-described markers of naïve and formative pluripotency and of the three germ layers confirmed elevated expression of naive ESC markers and reduced expression of formative and germ layer markers (Fig. 2c). Specifically, naïve-specific markers such as *Esrrb, Dppa4, Utf1*, and *Prdm14* were up, while formative markers such as *Fgf5* and *Dnmt3b* were down. Mesoderm (e.g. *Mixl1, Col4a1, Col4a2, Eomes*, and *T*), endoderm (e.g. *Sox17, Gata4, Hnf4a/b, Foxa2*), and ectoderm markers (e.g. *Pax6, Ncam1, Sox1* and *Krt18*) were consistently down (Fig. 2d,e). qRT-PCR analysis for a subset of these markers confirmed these findings.

**Figure 2.**
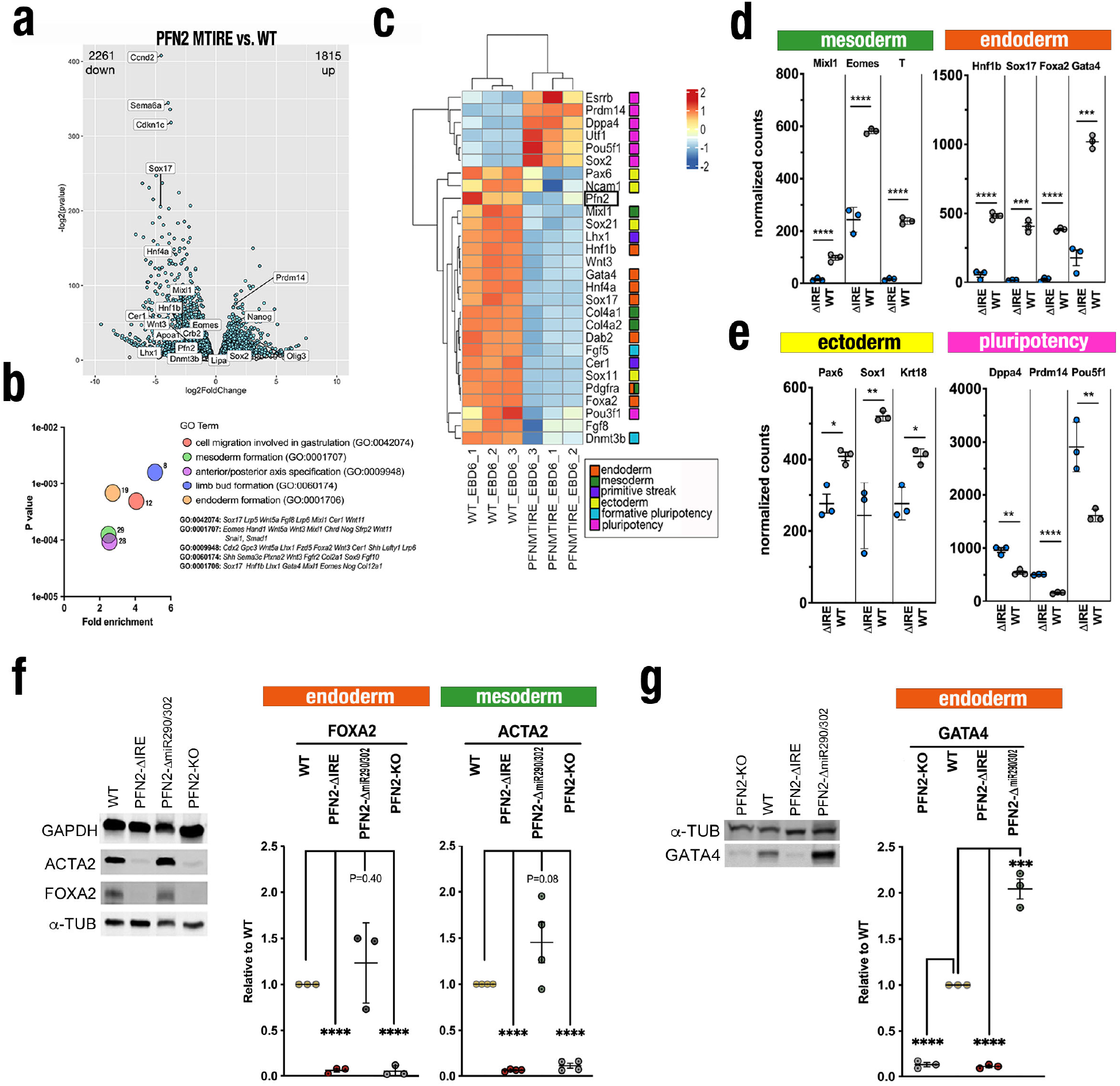
Deletion of IRE in PFN2-3UTR impairs early mesendodermal lineage specification. **(a)** Volcano plot of transcriptomic data PFN2-3UTR ΔIRE versus wild-type at EBD6 showing downregulated (left side) and upregulated (right side) differentially-expressed genes (FDR<0.05). Integration of n=3 biological replicates for each genotype across independent experiments. **(b)** Gene ontology enrichment analysis of PFN2-3UTR ΔIRE versus wild-type at EBD6. **(c)** Heatmap display showing mean-scaled normalized gene expression, measured by RNA-seq, of representative pluripotency and trilineage differentiation markers in wild-type and PFN2-3UTR ΔIRE mutants at EBD6; (bottom right) color coding scheme for known developmental and pluripotency markers **(d)** Dot plot of normalized counts of select representative transcripts of mesoderm, endoderm and **(e)** ectoderm, pluripotency-related markers from bulk transcriptomic studies of wild-type and PFN2-3UTR ΔIRE at EBD6. N=3 biological replicates for each genotype across independent experiments. Error bars represent SEM. *P< 0.05, **P< 0.01, ***P<0.001, ****P<0.0001, unpaired two-tailed t test. **(f-g)** Representative immunoblot (left) and immunoblot summary of protein levels (right) of mesoderm (ACTA2) and endoderm (FOXA2, GATA4) markers in wild-type versus PFN2-3UTR ΔIRE, PFN2-3UTR ΔmiR290/302, and PFN2-KO mutants at EBD6. Scanned images of unprocessed blots are shown in Fig. S2.

While the miR290/302 site mutants showed reduced expression of the ectoderm marker Pax6, there were normal levels of mesoderm (Mixl1) and endoderm (Gata4, Ttr) markers (Fig. S2). Immunoblot analysis showed that proteins associated with mesoderm (ACTA2) and endoderm (FOXA2, GATA4) were reduced in the IRE site mutant cells at EBD6 (Fig 2f-g). A similar phenotype was seen with *Pfn2* knockout cells. In contrast, abnormal upregulation of these proteins was seen in the miR290/302 site mutants. Together, these results show a striking defect in ESC differentiation with the loss of the IRE site in the *Pfn2* 3’UTR. In contrast, loss of the miR290/302 site appears to result in a defect in ectoderm differentiation consistent with the prior report of an FGF-related defect in the naïve to formative pluripotency differentiation (4). Therefore, these two 3’UTR elements have divergent impact on both PFN2 levels and cell fate during ESC differentiation.

### Loss of the PFN2-3UTR IRE site results in a cell autonomous defect in Wnt signaling

We noticed the PFN2-3UTR ΔIRE mutant EBs were morphologically abnormal compared to wild-type between EBD6 and EBD6.5, showing a loose dissociation at the outer surface of the EBs followed by failure to differentiate or cavitate (Fig. 3a-d). This dysmorphic defect appeared similar to a previously reported morphological defect associated with EB differentiation of ESCs deficient in *Ctnnb1* (beta-catenin) (33). Beta-catenin plays important roles in both cell adhesion through its interaction with cadherins at cell-cell junctions and in cell signaling as a secondary messenger in canonical Wnt signaling (33,34). Importantly, Wnt signaling is essential for mesendoderm differentiation during ESC differentiation (35). Given the combined PFN2-3UTR ΔIRE defects in EB morphogenesis, differentiation into mesoderm and endoderm lineages, we hypothesized a role for increased PFN2 in regulating beta-catenin and enabling differentiation of ESCs into the three germ layers.

**Figure 3.**
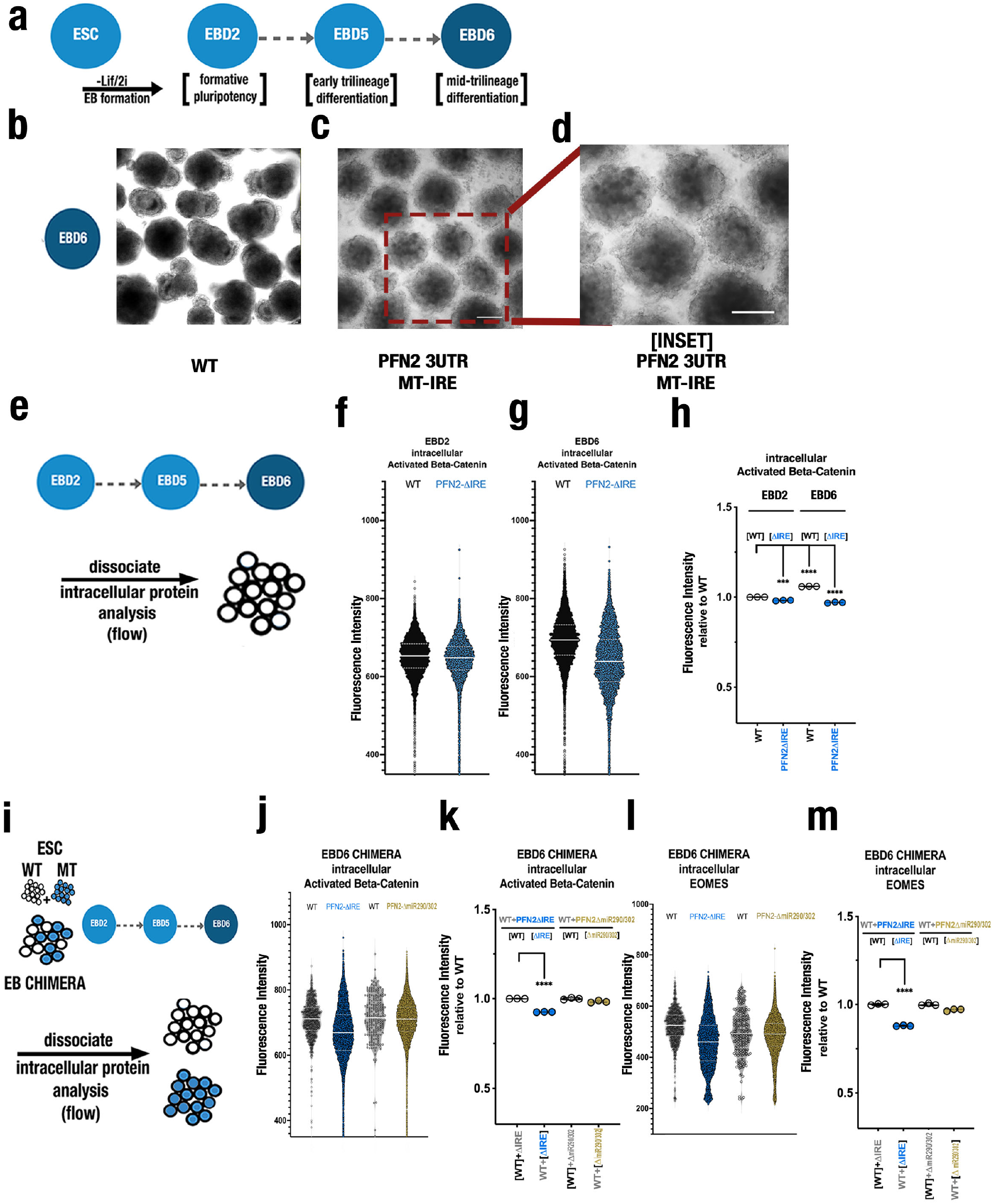
Deletion of IRE inPFN2-3UTR leads to morphologic and molecular defects in embryoid bodies during differentiation. **(a)** Schematic of experimental setup of stem cells (ESC) to embryoid body (EB) formation and differentiation (EBD2-6) with associated developmental trajectory. Representative images of **(b)** wild-type and **(c)** PFN2-3UTR ΔIRE mutant EBD6 embryoid bodies in culture, with **(d)** focus inset highlighting morphologic defect **(e)** Schematic of EB experimental culture and analysis process **(f)** Flow cytometry analysis of dissociated cells from wild-type versus PFN2-3UTR ΔIRE mutant embryoid bodies stained for active (non-phosphorylated) beta-catenin at **(f)** EBD2 and **(g)** EBD6, with **(h)** summary plot of n=3 independent experiments (single cells of representative experiment shown), ***P<0.001, ****P<0.0001, unpaired two-tailed t test. **(i)** Schematic depicting chimera formation and EB time course, intracellular protein analysis. Representative flow cytometry analysis of dissociated **(j)** wild-type versus PFN2-3UTR ΔIRE mutant chimeric embryoid bodies and wild-type versus PFN2-3UTR ΔmiR290/302 mutant chimeric embryoid bodies stained for intracellular active beta-catenin at EBD6, with associated **(k)** summary plot of n=3 independent experiments (single cells of representative experiment shown), ****P<0.0001, unpaired two-tailed t test. Representative flow cytometry analysis of dissociated **(l)** wild-type versus PFN2-3UTR ΔIRE mutant chimeric embryoid bodies and wild-type versus PFN2-3UTR ΔmiR290/302 mutant embryoid bodies stained for intracellular EOMES protein at EBD6, with associated **(m)** summary plot of n=3 independent experiments (single cells of representative experiment shown), ****P<0.0001, unpaired two-tailed t test.

Beta-catenin levels are regulated by GSK3β-directed phosphorylation, a post-translational modification that directs beta-catenin to the proteosome for degradation. In the absence of Wnt signaling, beta-catenin is associated with the destruction complex that includes sequential phosphorylation of Thr41, Ser37, and Ser33 by GSK3β (36). The presence of Wnt-signaling blocks this beta-catenin phosphorylation and degradation (37). Wnt/β-catenin signaling and subsequent β-catenin/TCF transcriptional activation are specifically mediated through the molecular form of β-catenin that remains unphosphorylated at residues 37 and 41 (38,39) often referred to as the signaling form of β-catenin, or Active β-Catenin (ABC). Therefore, to test our hypothesis, we performed analysis of intracellular unphosphorylated (active) beta catenin levels at single cell resolution by flow cytometry at EBD2 and at EBD6 (Fig. 3e), At EBD2, the intracellular levels of active beta-catenin in wild-type and PFN2-3UTR ΔIRE mutant EBs were not significantly different, while at EBD6, active beta-catenin levels were significantly lower in the PFN2-3UTR ΔIRE mutant cells. (Fig. 3f-h). In contrast, active beta-catenin levels were not impacted in PFN2-3UTR ΔmiR290/302 mutants at either EBD2 or EBD6 (Fig. S3a-c).

Next, we asked if the decrease in active beta-catenin was cell autonomous (e.g. defects in signaling downstream of the Wnt receptor or non-autonomous (e.g. defect in Wnt secretion). To do so, we produced chimeric EBs mixing mutant with wild-type cells expressing distinct color labels (Fig. S3d). Chimeric EBs composed of both wild-type and PFN2-3UTR ΔIRE showed less of a morphological phenotype than EBs solely made up of PFN2-3UTR ΔIRE cells, although their surfaces still appeared less sharp than WT+WT or WT+PFN2-3UTR ΔmiR290/302 chimeric EBs (Fig. S3e-g). Also, there was skewing of PFN2-3UTR ΔIRE cells toward the center of the EBs (Fig. S3h-j). Next, we dissociated the EBs into single cells and performed flow cytometric analysis for the color labels, intracellular beta-catenin, and intracellular EOMES as a marker of mesendoderm differentiation (Fig. 3i). This analysis showed that the PFN2-3UTR ΔIRE, but not wild-type or PFN2-3UTR ΔmiR290/302 cells isolated from the chimeras showed reduced active and total beta-catenin (Fig. 3j,k, Fig. S3k,l). Furthermore, unlike the wild-type or PFN2-3UTR ΔmiR290/302 cells, the PFN2-3UTR ΔIRE cells derived from the chimeras failed to upregulate EOMES, (Fig. 3l,m). Together, these data uncover a cell-autonomous defect in active beta-catenin signaling downstream of the Wnt receptor in the PFN2-3UTR ΔIRE background.

### PFN2-3UTR IRE site functions downstream of GSK3 to stabilize beta-catenin

To look further into mechanism by which PFN2 impacts beta-catenin and cell differentiation, we transitioned to a directed mesendodermal differentiation protocol (Fig. 4a). EBs were produced as above, but at day 3, CHIR and Activin were added to the media to specifically induce mesendoderm differentiation (35,40–43). T Brachyury (*Tbxt*) upregulation was used as the readout of successful differentiation into early mesendodermal progenitors (35,42). T Brachyury is also a direct target of Wnt induced beta-catenin signaling (44). Wild-type cells showed an over 30x increase in *Tbxt* (Fig. 4b). In contrast, PFN2-3UTR ΔIRE cells showed greatly reduced induction (∼2x). Intracellular flow cytometry of active beta-catenin remained reduced in PFN2-3UTR ΔIRE relative to wild-type cells even in the presence of the GSK3 inhibitor (Fig. 4c,d). To further confirm the cell autonomous nature of the defect, we repeated the chimera experiments, now under directed mesendodermal culture conditions. Similar to undirected EB differentiation, combining PFN2-3UTR ΔIRE and wild-type cells failed to rescue either active beta-catenin or total beta-catenin levels in directed differentiation conditions (Fig. 4e-g). By contrast, there were no significant differences between wild-type and PFN2-3UTR ΔmiR290/302 mutants within chimeras at EBD4M. Together, these results show that PFN2 is acting downstream of GSK3β directed beta-catenin degradation to regulate Wnt-signaling and thus mesendodermal differentiation.

**Figure 4.**
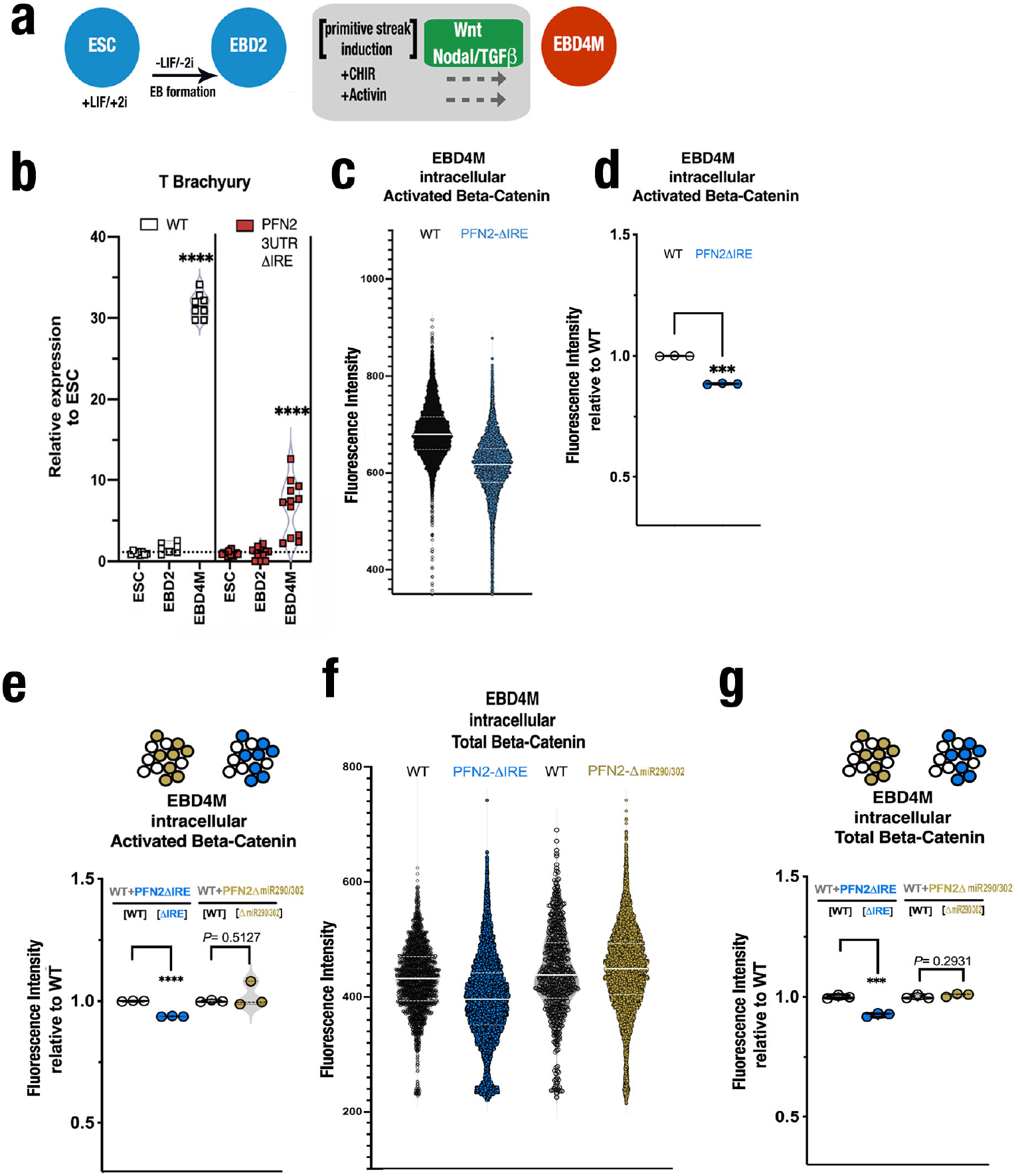
Deletion of IRE in PFN2-3UTR IRE impairs beta catenin levels and Brachyury expression during primitive streak-like (Wnt/Nodal) induction. (**a)** Schematic experimental setup for directed mesoderm differentiation. **(b)** Relative expression of Brachyury (T) in PFN2-3UTR ΔIRE mutants versus wild-type by quantitative RT-PCR at ESC, EBD2, and EBD4M. N= 3 biological replicates for each genotype across independent experiments. ESC vs EBD4M, **** = P<0.0001, unpaired two-tailed t test. **(c)** Representative flow cytometry analysis of one experiment of dissociated wild-type versus PFN2-3UTR ΔIRE mutant embryoid bodies stained for active (non-phosphorylated) beta-catenin at EBD4M, with associated **(d)** summary plot of n=3 independent experiments, ***P<0.001, unpaired two-tailed t test. Representative flow cytometry analysis of dissociated **(e)** summary plot of wild-type versus PFN2-3UTR ΔIRE mutant chimeric embryoid bodies (left) and wild-type versus PFN2-3UTR ΔmiR290/302 mutant chimeric embryoid bodies (right) stained for active (non-phosphorylated) beta-catenin at EBD4M, **(f)** wild-type versus PFN2-3UTR ΔIRE mutant chimeric embryoid bodies (left) and wild-type versus PFN2-3UTR ΔmiR290/302 mutant chimeric embryoid bodies (right) stained for total beta-catenin at EBD4M, (single cells of representative experiment shown), with **(g)** summary of multiple experiments, ***P<0.001, unpaired two-tailed t test.

### PFN2-3UTR IRE regulates nuclear beta-catenin levels

Wnt/beta-catenin signaling requires the transport of beta-catenin into the nucleus where it acts as a co-transcription factor in association with TCF/LEF proteins (Fig 5a). Given PFN2’s known association with the cytoskeletal network (45), we hypothesized that destabilization of beta-catenin could be secondary to a failure of beta-catenin transport to, retention of, or other mechanism of accumulation in the nucleus. To test this hypothesis, we performed a time-course of directed mesendoderm differentiation immediately following CHIR-Activin induction at EBD3, collecting samples at 2, 6, and 18 hours post-induction, and performing nuclear enrichment (Fig. 5b-g, Fig. S4). Nuclear active beta-catenin peaked at 18H in wild-type cells and was ∼3x fold reduced in PFN2-3UTR ΔIRE cells (Fig. 5d, Fig. S4). In contrast, cytoplasmic levels of either active or total beta-catenin were not significantly different between the wild-type and mutant cells (Fig. 5e-f). Nuclear T Brachyury levels similarly peaked at 16 hours and were dramatically reduced in PFN2-3UTR ΔIRE cells (Fig. 5g). ELISA of PFN2 itself showed reduced levels in both the cytoplasmic and nuclear compartments during the induction time course (Fig. 5h) and at the EBD4M endpoint (Fig. 5i). Together, these results indicate a possible defect in transport, retention, or accumulation of nuclear beta-catenin associated with the reduced levels of PFN2 in PFN2-3UTR ΔIRE cells.

**Figure 5.**
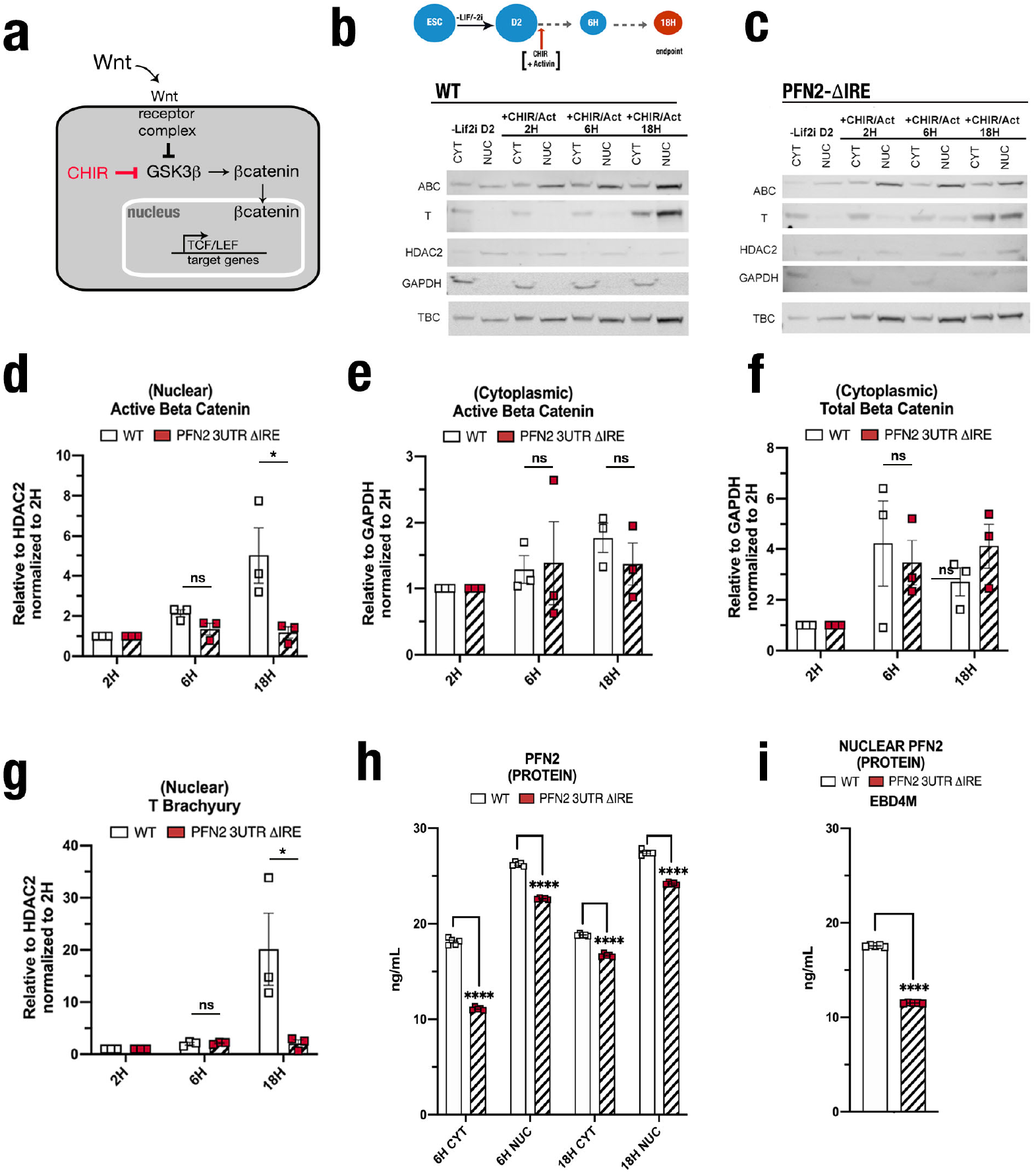
Deletion of IRE in PFN2-3UTR is associated with decreased Profilin2 levels, decreased nuclear retention of active beta-catenin, and failure to induce T Brachyury. **(a)** Schematic of beta-catenin signaling pathway showing including cytoplasmic-nuclear transit of beta-catenin following activation leading to TCF/LEF target gene transcription inducing mesendodermal differentiation. **(b)** Schematic experimental setup for time course collection following addition of CHIR+Activin.(top) Immunoblot analysis of active beta-catenin (ABC), total beta-catenin (TBC), and Brachyury (T) protein levels in cytoplasmic and nuclear subcellular compartments in wild-type (bottom) and **(c)** PFN2-3UTR ΔIRE mutants over 18 hour differentiation timecourse. Scanned images of unprocessed blots are shown in Fig. S4. Protein levels of **(d)** nuclear active beta-catenin, **(e)** cytoplasmic active beta-catenin, **(f)** cytoplasmic total beta-catenin, and **(g)** nuclear Brachyury (T), and in wildtype (white) and PFN2-3UTR ΔIRE mutants (red) over 18 hour timecourse. **(h)** Subcellular cytoplasmic and nuclear PFN2 protein by ELISA in wild-type (white) and PFN2-3UTR ΔIRE mutants (red) at 6H and 18H timepoints. **(i)** Nuclear PFN2 protein by ELISA in EBD4 lysate from wild-type (white) and PFN2-3UTR ΔIRE mutants (red). (h-m) N=3. Error bars represent SEM. *P< 0.05, ****P<0.0001, unpaired two-tailed t test.

## Discussion

Together with the recent dissection of the miR290/302 site in the 3’UTR of PFN2 (4), the findings reported here uncover a coordinate role of a microRNA and an RNA binding protein on the 3’UTR of a single gene in regulating the switch from pluripotency to somatic differentiation (Fig. 6). In pluripotent cells, the dominant miR-290 cluster destabilizes PFN2, allowing endocytosis which is essential for normal FGF/ERK signaling (4). This signaling is important for the normal transition from naïve to formative pluripotency (4,46–49). Following the transition, an IRE which is bound by iron response proteins is required for an increase in PFN2 levels, in turn promoting Wnt/beta-catenin signaling. Defects in the Wnt/beta-catenin due to deletion of the IRE result in a block in activation of the mesendoderm transcriptional program that normally occurs with gastrulation. The impact of PFN2 on beta-catenin signaling is downstream of GSK driven degradation of the protein and is associated with alterations in the nuclear/cytoplasmic localization of the protein suggesting a role in the transport, retention, and/or accumulation of beta-catenin in the nucleus. Thus, our findings show how the combinatorial post-transcriptional control of PFN2 plays a critical role in early mammalian cell fate decisions.

**Figure 6.**
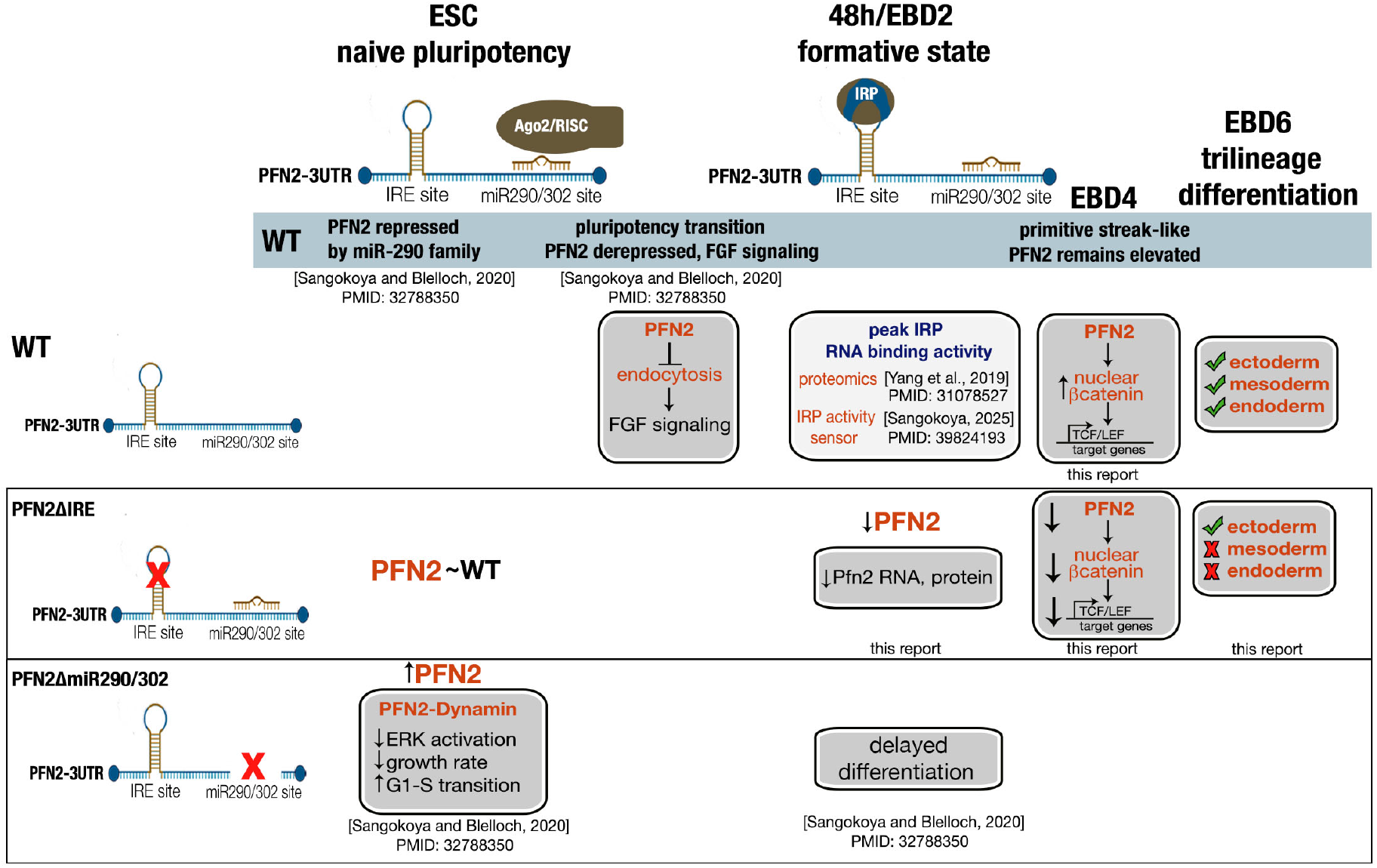
Impact of post-transcriptional control of the Profilin-2 transcript on stem cell biology during pluripotency, pluripotency transition, and early trilineage differentiation. Schematic summary of the impact of Profilin-2 transcript Ago2/miR-290 and IRP/IRE binding sites on stem cell biology during pluripotency, pluripotency transition, and early trilineage differentiation.

Previous examples of microRNA and RBP coordination on a 3’UTR largely involve RBPs enhancing microRNA targeting by improving target site accessibility (50–52) or suppressing microRNA targeting by competitive engagement of overlapping binding sites (52–57). In the setting of development, for example, the RNA-binding protein Dead end (Dnd1) prohibits the function of several miR-430 and miR-372-family microRNAs by blocking their accessibility to transcript target sites, and this regulation is essential for proper development of primordial germ cells (53). In the case of PFN2, the RBP and microRNA binding sites are not overlapping, but instead are ∼100 bases apart and are acting at distinct developmental times. Therefore, it seems unlikely that their opposing impacts on PFN2 levels are due to competitive binding. Instead, these sites reveal a developmentally relevant switch where the predominance of miR-290 microRNA regulation on PFN2 during pluripotency ‘switches’ to a predominant role for IRP binding during somatic differentiation into the three germ layers. We propose that this model could provide a paradigm for other microRNA-RBP pairs playing critical roles in cell fate decisions throughout development.

Our data further shows that PFN2 regulates the transport, retention, or accumulation of beta-catenin in the nucleus. Reduced PFN2 levels during ESC differentiation by deletion of the IRE in its 3’UTR results in reduced nuclear beta-catenin levels without significantly impacting cytoplasmic levels. Nuclear beta-catenin is critical for the differentiation of embryonic stem cells down the meso-endoderm lineages (35). Therefore PFN2’s role in regulating nuclear beta-catenin levels can explain the requirement for elevated PFN2 levels during ESC differentiation into mesendoderm. These findings raise the important question of how PFN2 regulates beta-catenin nuclear import and/or retention. A small number of beta-catenin interacting proteins have been shown to play a role in its nuclear import, but otherwise very little is known (58–60). Our attempts at co-immunoprecipitation of PFN2 and beta-catenin failed to uncover a direct interaction (data not shown). Therefore, it remains unclear if the role of PFN2 in beta-catenin cellular localization is direct or indirect. However both PFN2 and beta-catenin interact with actin (61– 63) and, therefore, it could be that a direct role for PFN2 in actin control is indirectly impacting beta-catenin nuclear import and/or retention. Indeed, nuclear transport of beta catenin and actin have been proposed to be tightly intertwined (64). Beta-catenin is a member of the Armadillo family, whose members share the ability to move into the nucleus independent of the classic nuclear localization signal (NLS) which uses an Importin-dependent mechanism for transport (65). Therefore, it is possible that PFN2 will be found to play important roles in the transport of other Armadillo families as well. In either case, future studies on PFN2’s role in beta-catenin nuclear transport/retention or accumulation are likely identify further novel insights into how the Wnt pathway is controlled in time and space during development.

In summary, our findings reveal that post-transcriptional regulation of the cytoskeletal protein Profilin-2 within a conserved short region of its 3’UTR that includes both an Ago2/microRNA site and an IRP/IRE site is essential for its tight molecular control during early ESC differentiation. This tight control of Profilin-2 expression tunes the FGF/ERK signaling pathway, which is integral to the naïve to formative pluripotency transition, as well as promotes the WNT/Beta-catenin signaling pathway, which is integral to lineage commitment to mesendoderm. This model provides a paradigm for how the coordination of microRNA and RBP binding control on a single transcript over development time can regulate cell fate decisions.

### Limitations of study

While this study highlights a previously undescribed developmental switch based on post-transcriptional regulation, there are limitations and unanswered questions. (1) It remains unresolved whether and which extended protein interaction networks interact with PFN2 to modulate nuclear beta-catenin in the context of ESC differentiation. (2) This study is unable to distinguish precisely how PFN2 regulates nuclear beta-catenin – whether by transport, retention, or action on mechanisms directly regulating nuclear accumulation. (3) Of note, in the PFN2-3UTR ΔIRE mutant cells, we appreciated an unexplained slight increase in Pfn2 protein levels at EBD2 in Fig. 1e, although still significantly depleted compared to wild-type cells. Other isoforms of the mouse Pfn2 mRNA have been described (66,67) that differ in the last exon and yield protein with different C-terminal amino-acid composition which do not retain the 3′-IRE. Despite these limitations, this study reveals unexplored areas for further investigation. Future studies utilizing parallel and intersectional transacting and protein-based approaches to illuminate the spatial context and signaling networks during ESC differentiation would therefore be useful to confirm and extend these findings.

## Acknowledgments

Flow Cytometry was performed at the UCSF Parnassus Flow Cytometry Core, supported in part by the DRC Center Grant NIH P30 DK063720. Sequencing was performed at the UCSF Center for Advanced Technology, supported by UCSF PBBR, RRP IMIA, and NIH 1S10OD028511-01 grants. This research was supported by funding from NIH/NICHD (K08HD105017) and Burroughs Wellcome Fund (to C.S.), and R01 GM125089 and GM122439 (to R.B.).

## Author Contributions

C.S. and R.B. developed the concept and designed experiments. C.S. conducted all experiments and all data analysis. C.S. and R.B. interpreted the data, wrote and edited the manuscript.

## Methods

### Cell Culture and ESC Monolayer Differentiation

Mouse V6.5 ESCs (RRID:CVCL_C865) were grown and maintained at 37C and 5%CO2 on 0.2% gelatin-coated plates, cultured in ESC medium (custom DMEM) supplemented with 15% FBS (Corning Mediatech), LIF (1000U/mL, custom) and 2i (1uM PD0325901 (Axon MedChem) and 3uM CHIR99021 (Axon MedChem). Differential states of pluripotency from naïve to EpiLC transition were generated by removal of LIF and 2i. In brief, ESCs were plated in ESC medium (as described above). To initiate differentiation, LIF and 2i were removed. Cells were collected at indicated time points following LIF/2i removal.

### Embryoid Body Generation and Differentiation

For embryoid body (EB) generation, ESCs were cultured in rotary (100 rpm) suspension culture on a low attachment plate allowing for self-aggregation and regular reproducible spontaneous differentiation over a course of 7 to 10 days. EB differentiation was evaluated by quantitative reverse transcriptase PCR, immunoblot analysis, and bulk RNA-sequencing. Directed differentiation to mesendoderm was performed by induction with 10 ng mL^-1^ Activin (GF300, Millipore) and 3 uM CHIR99021 (Axon MedChem). Chimeric embryoid bodies were formed by combining equal numbers of cells prior to rotary suspension culture.

### Quantitative RT -PCR

RNA was extracted from cells using TRIzol (Invitrogen) and quantified on a Nanodrop spectrophotometer (ThermoFisher Scientific). Reverse transcription and quantitative PCR were performed according to manufacturer’s instructions (Maxima, K1641, ThermoScientific; PowerSYBR, Applied Biosystems). Primer sequences are listed in [Supplementary Table 1].

### ELISA immunoassay

Sandwich ELISA immunoassay for in vitro quantitative measurement of PFN2. A microplate pre-coated with an antibody specific for Mouse PFN2 was used for sandwich ELISA immunoassay in vitro quantitative measurement of PFN2 (RK06647, ABclonal Science) according to manufacturer’s instructions.

### Generation of Site-based Mutant Cell Lines by CRISPR-CAS9 Gene Editing

Mutant ESC cell lines were generated by cloning gRNAs into a U6 vector (GE100042, Origene) and co-transfection with a Cas9-expressing vector (GE100018, Origene) using Fugene6 (E2691, Roche) as described in (4). Guide sequences are listed in [Supplementary Table 2].

### RNA-seq library preparation and sequencing

RNA (total or poly-adenylated) was extracted using TRIzol (15596026, Invitrogen) or Oligo(dT)25-coated Dynabeads (61002, Invitrogen), respectively. RNA quality was examined by Qubit 2.0 Fluorometer (Life Technologies). cDNA library generation was done using the KAPA RNA Hyperprep kit (KK8541, Roche) following the manufacturer’s instructions. cDNA library quality was assessed using Qubit 2.0 fluorometer to determine concentrations. The 4200 TapeStation system (Agilent) was used to determine insert size following the manufacturer’s instructions. cDNA libraries were then sequenced using the HiSeq 4000 (Illumina) and NovaSeq X Systems (Illumina).

### Bulk RNA-Seq data pre-processing and analysis

Raw fastq files were filtered and preprocessed by quality trimming (-Q33, threshold 20, minimum length 15) using FASTX-Toolkit (http://hannonlab.cshl.edu/fastx_toolkit, (68)). Adapters were removed and reads collapsed using unique molecular identifiers using FASTX-Toolkit and Cutadapt (56). RNA-seq read mapping and alignment with alignment to mm10 reference genome assembly was performed using STAR (69). Read count tables were generated using the RSEM program (70). Differential gene expression was performed using R package DESeq2 (71) for analysis including plotCounts to generate normalized plot counts. R (72) packages including edgeR (73), pheatmap (74) were used for data visualization. FDR was calculated using the Benjamini–Hochberg method. DE genes were defined with FDR less than 0.05 in comparison to the control group. Default settings were used unless otherwise specified.

### Gene Ontology and GO Enrichment Analysis

Gene Ontology reference and GO Enrichment Analysis were accessed and used to perform ontology and fold enrichment analysis using the GO Ontology database DOI: 10.5281/zenodo.10536401 (75,76). Differentially expressed transcripts (with p<0.05, FDR <0.05, and log fold change greater than an absolute value of 0.06, for at least a 1.5 fold cutoff) were uploaded to the Gene Ontology Resource at https://geneontology.org. From there, the analysis tool from the PANTHER Classification System (76) is used to perform enrichment analyses by biological process. Results with threshold FDR<0.05 are reported.

### Immunoblot

Total cell extracts were obtained by lysing cell pellets with ice-cold RIPA lysis buffer containing EDTA-free protease inhibitor mixture (11836170001, Roche) and PhosSTOP phosphatase inhibitor (4906837001, Roche), with DNA shearing by 21-gauge needle. Lysates were incubated at 4C for 15 min and centrifuged at 20,000 x g at 4 C on a table-top centrifuge. The protein concentration of the supernatant was measured by BCA assay, with 30 micrograms of protein resolved by sodium dodecyl sulfate-polyacrylamide gel electrophoresis. After electrophoresis, proteins were transferred onto an Immobilon-FL PVDF membrane (IPFL00010, Millipore) and processed for immunodetection. Primary antibodies were incubated overnight at 4C. Secondary antibodies were incubated for 1h at RT. Images were digitally acquired using a Li-COR Odyssey imager (Li-COR Instruments) and its proprietary software. Antibodies are listed in Supplementary Table 3. Bands were quantified using Image J (77).

### Subcellular Fractionations

Subcellular fractionations were performed according to manufacturer instructions (78840, Thermo Scientific), and followed by ELISA and/or immunoblot.

### Intracellular Staining Assay

Cells were first fixed with 4% paraformaldehyde (J61899, ThermoScientific) for 15 min at room temperature, then permeabilized on ice with 0.1% Triton-X (A16046.AE, ThermoScientific) for 5 min and stained for the indicated primary antibody for 30 min at 4C at dilutions as listed in Supplementary Table 3. This was followed by incubation for 30 minutes at 4C with species-specific secondary antibodies at dilutions as listed in Supplementary Table 3. Finally the cells were washed with cold PBS and prepared for analysis by flow cytometry. Antibodies are listed in Supplementary Table 3.

### Flow Cytometry

After dissociation into single-cell suspension using Trypsin (25200072, ThermoFisher Scientific) or TrypLE (12604039, ThermoFisher Scientific), cells were resuspended in HBSS/1% BSA (14175103, ThermoFisher Scientific; A30075, RPI) and filtered for size by cell strainer (35 micron mesh). Cells were processed by flow cytometry on an LSR II flow cytometer (Becton Dickinson), with events collected by FACS Diva software (BD Biosciences) using a hierarchical gating strategy selecting for a homogenous cell population (forward-scatter area vs. side-scatter area) of single cells (forward-scatter width vs. forward-scatter area). Dot plots and mean fluorescence intensities for gated events were generated using FlowJo analysis software.

### Statistical Analysis

No data were excluded from the analyses. Statistical tests were performed in GraphPad Prism (v10). Data collection and analysis were not performed blind to the conditions of the experiments, but data analyses have been performed with identical parameters and software. Data representation and statistical analyses were also performed using R software. The number of biological replicates and independent experiments, both equal to or greater than 3, is indicated in figure legends. The statistical tests used are indicated in the figure legends.

### Authentication and additional information

V6.5 ES cells and mutant V6.5 ES cells (PFN2-3UTR mutants, PFN2-KO) were authenticated by genotyping (PCR and Sanger sequencing) and by undirected and directed differentiations described in this manuscript. Key experiments were performed within 10 passages under sterile conditions. All cell lines are tested for mycoplasma before use and annually.

## Supplemental information

Document S1. Figures S1-S4.

## Data availability

Sequencing data that support the findings of this study have been deposited in the Gene Expression Omnibus and are publicly available as of the date of publication. Source data and data supporting the findings of this study are deposited on Zenodo and publicly available as of the date of publication. Any other data supporting the findings of this study are available from the corresponding author upon reasonable request.

## Code availability

This paper does not report original code.

