## SupplementalFiles for "Coordinate post-transcriptional regulation by microRNAs and RNA binding proteins is critical for early embryonic cell fate decisions"

### Figure S1

**CRISPR editing for PFN2 IRE mutant generation**

**(a)** Schematic representation of CRISPR/Cas9-mediated gene editing strategy to generate Pfn2 mutant ESC lines. **(b)** Schematic highlighting guide RNA sequences used to generate CRISPR-based ESC mutant cell line PFN2-3UTR ΔIRE and sequenced deletion result for three clones compared to wild-type sequence. PFN2-3UTR ΔIRE clones assayed for Pfn2 RNA expression (result shown in Fig. 1d).

### Figure S2

**PFN2 3UTR mutant site effects on ESC trilineage differentiation**

**(a)** Relative expression by quantitative PCR of lineage markers in PFN2-3UTR ΔIRE and PFN2-3UTR ΔmiR290/302 mutants versus wild-type at EBD5 representing ectoderm (Pax6) mesoderm (Mixl1), and endoderm (Ttr, Gata4) **P< 0.01, ***P<0.001, ****P<0.0001, unpaired two-tailed t test. **(b)** Scanned images of unprocessed blots included in immunoblot summary of protein levels of mesoderm (ACTA2) and endoderm (FOXA2) markers in wild-type versus PFN2-3UTR ΔIRE, PFN2-3UTR ΔmiR290/302, and PFN2-KO mutants at EBD6 shown in Fig. 2f-g.

### Figure S3

**Decreased beta-catenin and impaired mesoderm specification are cell autonomous defects in cells with disrupted PFN2-3UTR IRE site**

**(a)** Representative flow cytometry analysis of one experiment of dissociated wild-type versus PFN2-3UTR ΔmiR290/302 mutants embryoid bodies stained for active (non-phosphorylated) beta-catenin at EBD2 and **(b)** EBD6, with **(c)** summary plot of n=3 independent experiments. ***P<0.001, ****P<0.0001, unpaired two-tailed t test. **(d)** Schematic depicting EB chimera formation. Brightfield image of chimeric embryoid bodies composed of **(e)** wild-type + wild-type cells, **(f)** wild-type + PFN2-3UTR ΔIRE cells, and **(g)** wild-type + PFN2-3UTR ΔmiR290/302 cells at EBD5. Composite fluorescent image (wild-type in blue, other cell type in red) of chimeric embryoid bodies composed of **(h)** wild-type + wild-type cells, **(i)** wild-type + PFN2-3UTR ΔIRE cells, and **(j)** wild-type + PFN2-3UTR ΔmiR290/302 cells at EBD5. **(k)** Representative flow cytometry analysis of dissociated wild-type versus PFN2-3UTR ΔIRE mutant embryoid bodies and wild-type versus PFN2-3UTR ΔmiR290/302 mutant embryoid bodies stained for intracellular total beta-catenin at EBD6, with associated **(l)** summary plot of n=3 independent experiments.****P<0.0001, unpaired two-tailed t test.

### Figure S4

**PFN2-3UTR IRE site disruption is associated with decreased Profilin2 levels, decreased nuclear retention of active beta-catenin, and failure to induce T Brachyury**

Scanned images of unprocessed blots for Figures 5b and 5c immunoblots (Blot A shown in main figures) and replicates (Replicate B and C, summarized in Fig. 5d-g).

### Figure S1

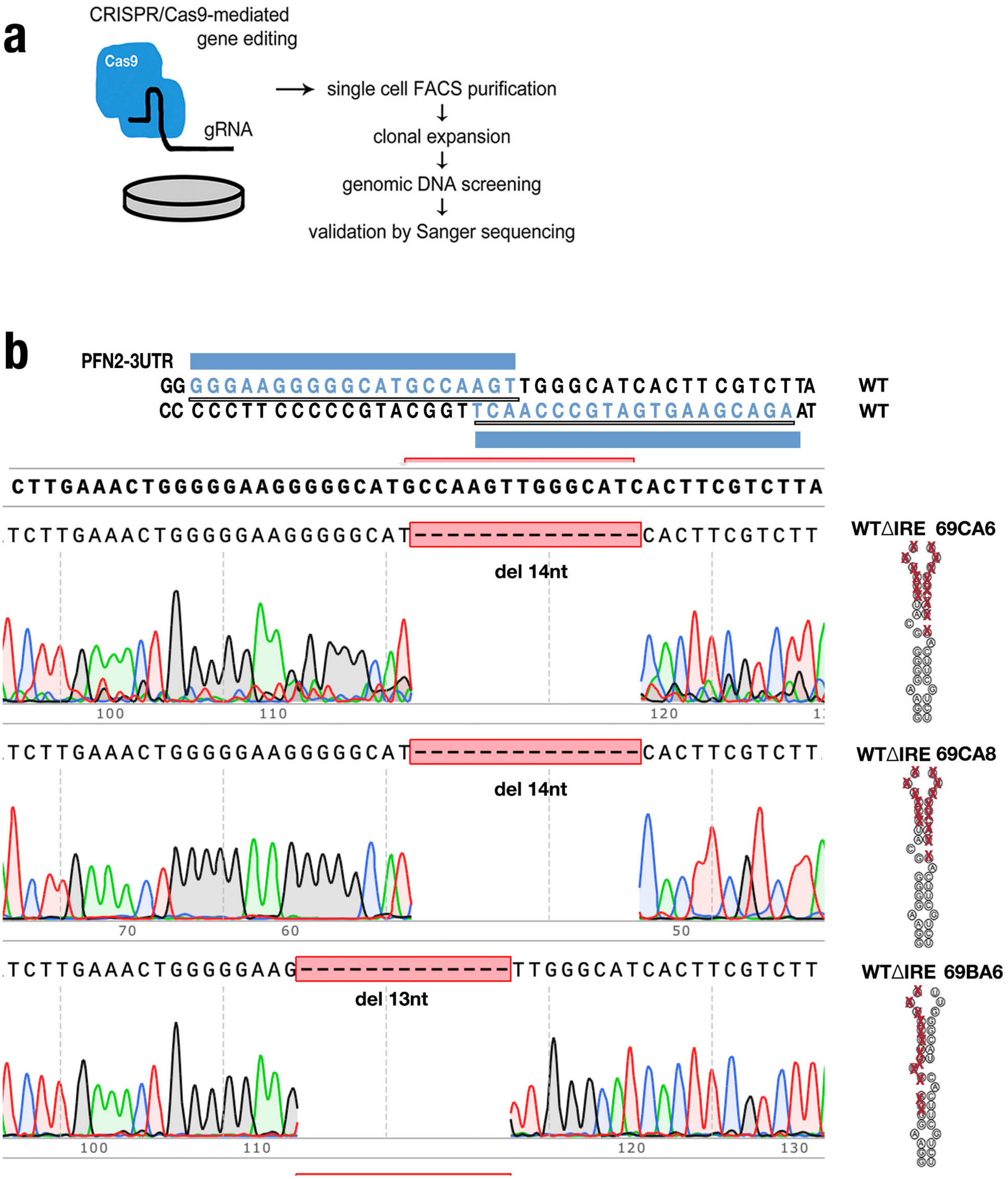
a
CRISPR/Cas9-mediated gene editing
Cas9
gRNA
single cell FACS purification
clonal expansion
genomic DNA screening
validation by Sanger sequencing
b
PFN2-3UTR
GGGGGAAGGGGGCATGCCAAGTTGGGCATCACTTCGTCTTA
WT
CCCCCTTCCCCCGTACGGTTCAACCCGTAGTGAAGCAGAAT
WT
CTTGAAACTGGGGGAAGGGGGCATGCCAAGTTGGGCATCACTTCGTCTTA
TCTTGAAACTGGGGGAAGGGGGCAT--------------CACTTCGTCTT
del 14nt
WTΔIRE 69CA6
100
110
120
TCTTGAAACTGGGGGAAGGGGGCAT--------------CACTTCGTCTT
del 14nt
WTΔIRE 69CA8
70
60
50
TCTTGAAACTGGGGGAAG-------------TTGGGCATCACTTCGTCTT
del 13nt
WTΔIRE 69BA6
100
110
120
130

### Figure S2

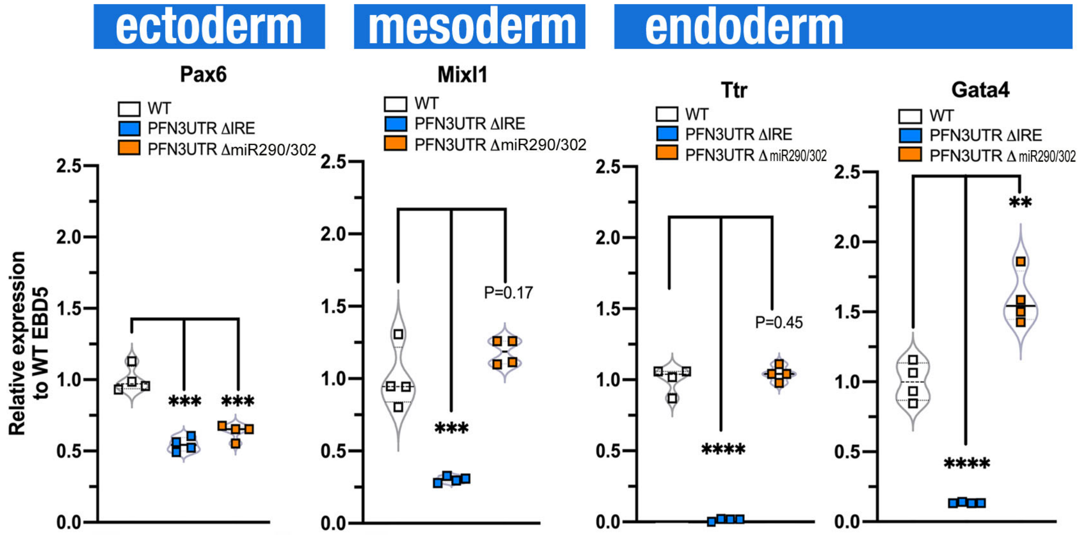


**Fig. 2f** n=4 shown on Blot A, Replicate B, Replicate C, Replicate D

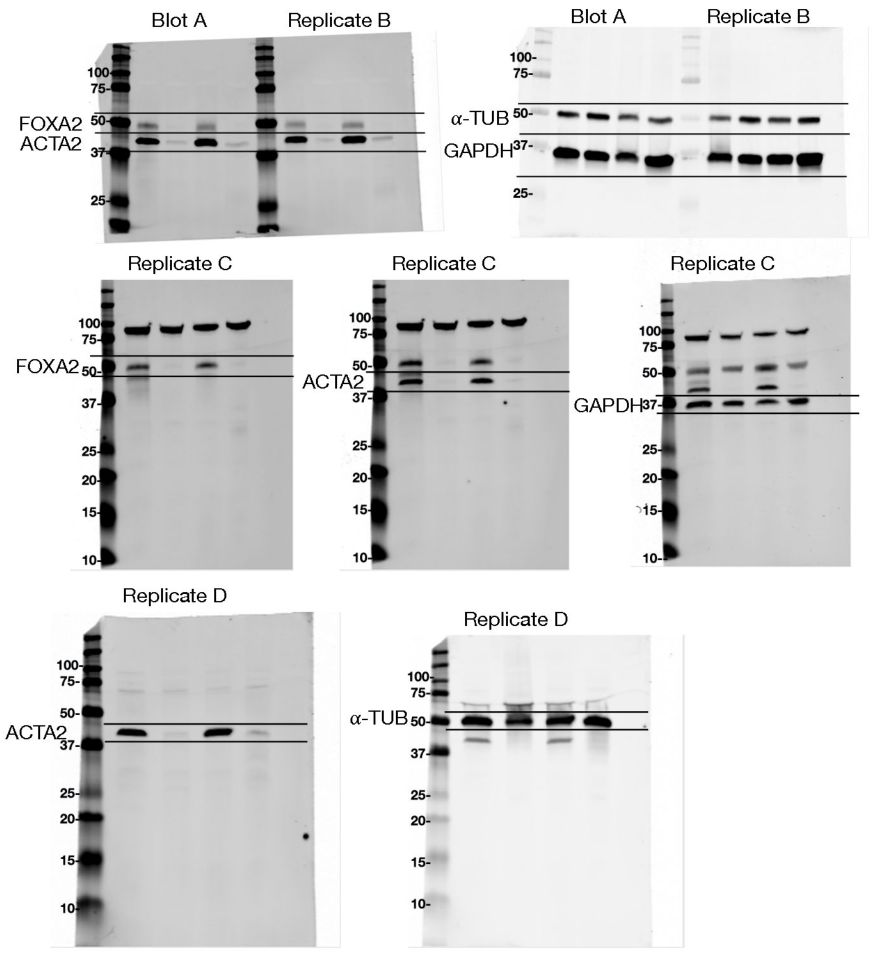


**Fig. 2g** n=3 shown on Blot A, Replicate B, Replicate C

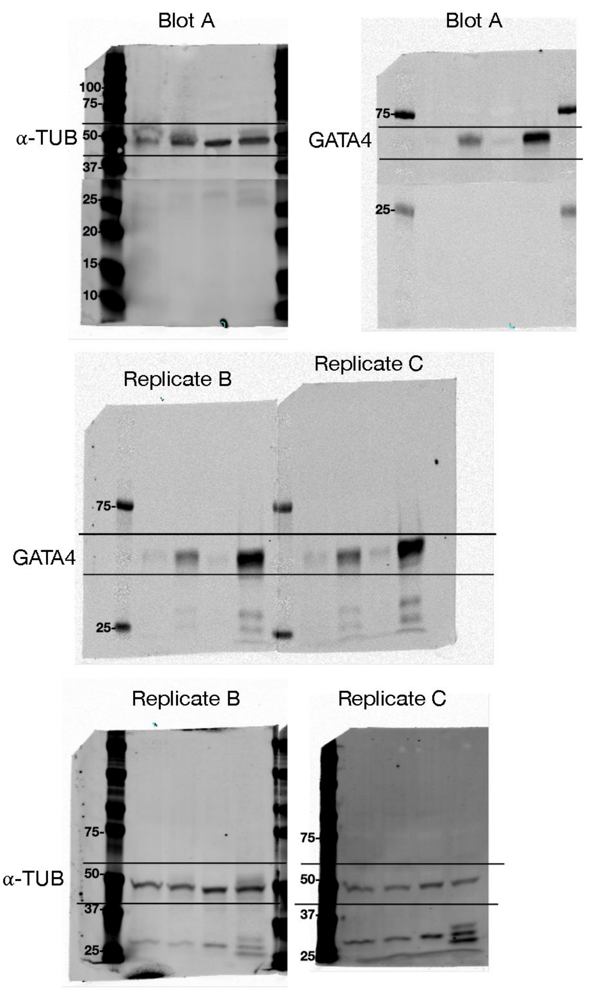

### Figure S3

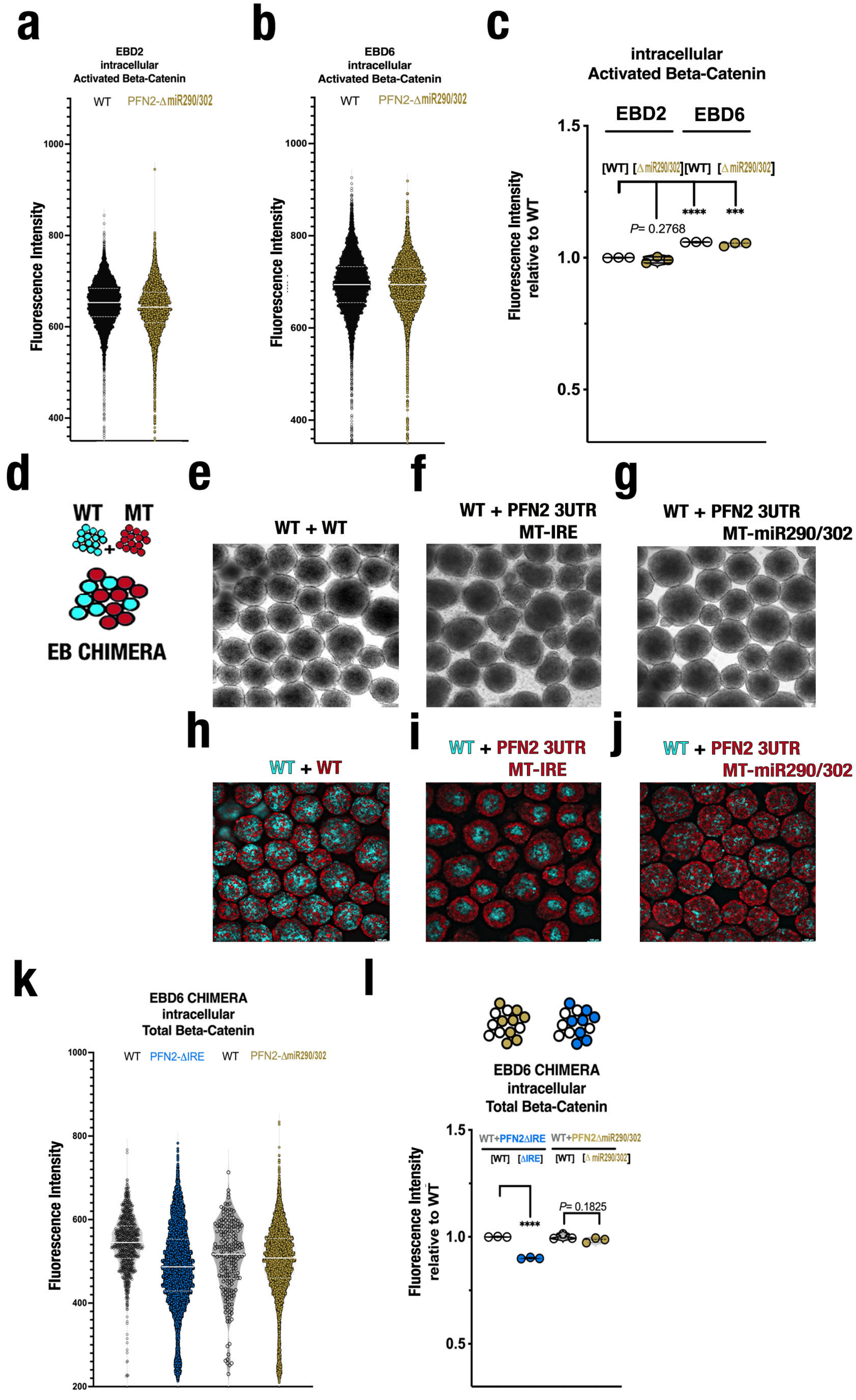

a
EBD2
intracellular
Activated Beta-Catenin
WT
PFN2-ΔmiR290/302
Fluorescence Intensity
1000
800
600
400
b
EBD6
intracellular
Activated Beta-Catenin
WT
PFN2-ΔmiR290/302
Fluorescence Intensity
1000
800
600
400
c
intracellular
Activated Beta-Catenin
EBD2
EBD6
[WT] [ΔmiR290/302] [WT] [ΔmiR290/302]
****
***
P= 0.2768
Fluorescence Intensity
relative to WT
1.5
1.0
0.5
d
WT
MT
EB CHIMERA
e
WT + WT
f
WT + PFN2 3UTR
MT-IRE
g
WT + PFN2 3UTR
MT-miR290/302
h
WT + WT
i
WT + PFN2 3UTR
MT-IRE
j
WT + PFN2 3UTR
MT-miR290/302
k
EBD6 CHIMERA
intracellular
Total Beta-Catenin
WT
PFN2-ΔIRE
WT
PFN2-ΔmiR290/302
Fluorescence Intensity
1000
800
600
400
200
l
EBD6 CHIMERA
intracellular
Total Beta-Catenin
WT+PFN2ΔIRE
WT+PFN2ΔmiR290/302
[WT] [ΔIRE]
[WT] [ΔmiR290/302]
****
P= 0.1825
Fluorescence Intensity
relative to WT
1.5
1.0
0.5

### Figure S4

#### Fig. 5b

C= CYT
N=NUC

0h= -LIF2i D2
2h= -LIF2i D2 +CHIR/Act 2h
6h=-LIF2i D2 +CHIR/Act 6h
18h=-LIF2i D2 +CHIR/Act 18h

n=3 shown on Blot A,
Replicate B, Replicate C

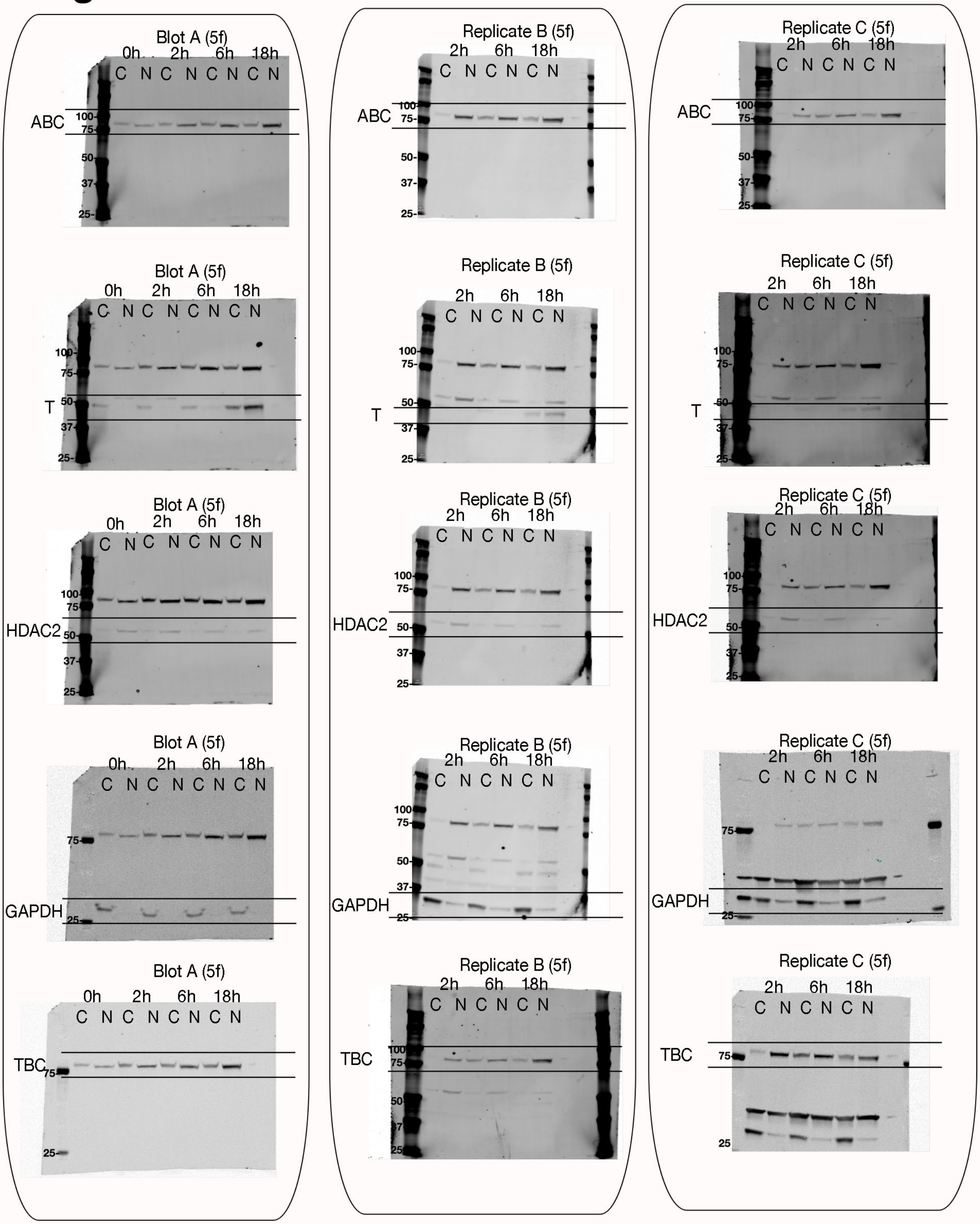

### Figure S4

#### Fig. 5c

C= CYT
N=NUC

0h= -LIF2i D2
2h= -LIF2i D2 +CHIR/Act 2h
6h=-LIF2i D2 +CHIR/Act 6h
18h=-LIF2i D2 +CHIR/Act 18h

n=3 shown on Blot A,
Replicate B, Replicate C

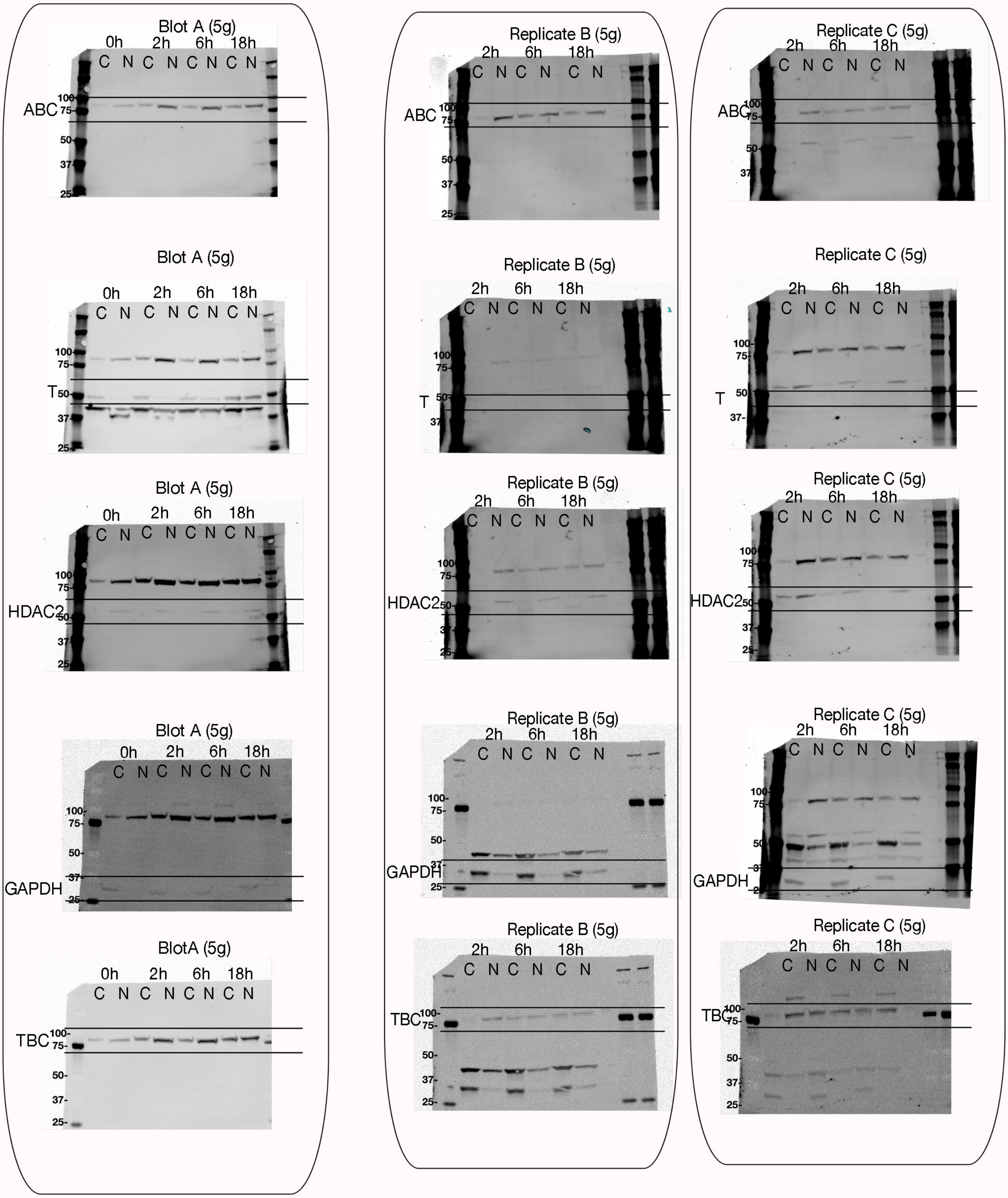
